# Evolutionary sweeps of subviral parasites and their phage host bring unique parasite variants and disappearance of a phage CRISPR-Cas system

**DOI:** 10.1101/2021.10.07.463549

**Authors:** Angus Angermeyer, Stephanie G. Hays, Maria H.T. Nguyen, Fatema-tuz Johura, Marzia Sultana, Munirul Alam, Kimberley D. Seed

## Abstract

*Vibrio cholerae* is a significant threat to global public health in part due to its propensity for large-scale evolutionary sweeps where lineages emerge and are replaced. These sweeps may originate from the Bay of Bengal where bacteriophage predation and the evolution of anti-phage counter defenses is a recurring theme. The bacteriophage ICP1 is a key predator of epidemic *V. cholerae* and is notable for acquiring a CRISPR-Cas system to combat PLE, a defensive subviral parasite encoded by its *V. cholerae* host. Here we describe the discovery of five previously unknown PLE variants, including one found during recent surveillance of patient samples in Bangladesh. We also observed a lineage sweep of PLE negative *V. cholerae* occurring within a patient population in under a year which coincided with a loss of ICP1’s CRISPR-Cas system. These findings reinforce the importance of surveillance to better understand the selective pressures that drive pandemic cholera.

## Introduction

Pandemics of the severe intestinal disease cholera have plagued humanity on a global scale for over two centuries. Caused by the bacterium *Vibrio cholerae*, the first pandemic is thought to have occurred in the early-mid 19^th^ century originating near the Bay of Bengal and sweeping across the Indian subcontinent, Asia and parts of east Africa (Sack et al., 2004). There have been six subsequent pandemics costing millions of lives and culminating in the current 7^th^ cholera pandemic which accounts for ∼120,000 deaths per year globally (Harris et al., 2012) and has spread heavily throughout Africa and Asia. Devastating outbreaks have also occurred in the Americas, Europe, Oceania and the Middle East. The genetic history of the 7^th^ pandemic is marked by several temporally overlapping waves of distinct *V. cholerae* lineages which themselves are comprised of numerous inferred transmission events (Weill et al., 2017). A hallmark of *V. cholerae’s* evolution is that not only do new lineages frequently arise, but that prior lineages simultaneously disappear. Pandemic *V. cholerae* is principally the O1 serogroup which is divided into ‘classical’ and ‘El Tor’ biotypes (Kaper et al., 1995). A particularly dramatic example of lineage replacement in *V. cholerae* is that the first six pandemics are thought to have been entirely O1/classical strains, while the 7^th^ pandemic is predominantly O1/El Tor (Cvjetanovic and Barua, 1972).

Isolates from within the waves of the current pandemic are highly homogenous and each can be traced back to the Bay of Bengal as the original source (Mutreja et al., 2011). Cholera outbreaks are endemic to communities surrounding this area where it is thought that *V. cholerae* is in continuous circulation between the human population and estuarian environments that act as the natural reservoir (Alam et al., 2006; Stine et al., 2008). *V. cholerae* is thought to undergo the majority of its diversification and selection in the aquatic reservoir, giving rise to novel lineages that can infect local communities, spread to cities and from there spread globally (Mavian et al., 2020). Due to this single-point origin of epidemic *V. cholerae*, surveillance of cholera along the coast of the Bay of Bengal, specifically in Bangladesh, has previously proven very important in identifying not only novel bacterial genotypes (Seed et al., 2013), but also factors in *V. cholerae’s* ecosystem that influence the competition between lineages and help shape the selection of genotypes that cause outbreaks (Blokesch and Schoolnik, 2007; Faruque and Mekalanos, 2014). Of these factors, bacteriophage predation is likely of particular importance given that phages specific to *V. cholerae* are found in water sources in endemic regions as well as in cholera patient stool (d’Herelle, 1961; Faruque and Mekalanos, 2014; Seed et al., 2011). With phage predation threatening *V. cholerae* throughout its lifecycle, the fitness of epidemic strains is likely contingent on the evolution of phage defense mechanisms.

Of the bacteriophages known to target *V. cholerae*, ICP1 stands out for its persistence having first been detected in 1992 in India (Boyd et al., 2021) and consistently recovered in stool samples from Bangladesh since then (Angermeyer et al., 2018; Boyd et al., 2021; McKitterick et al., 2019b; Seed et al., 2011). However, *V. cholerae* can defend against ICP1 through the acquisition of parasitic anti-phage islands called PLEs (<u>P</u>hage inducible chromosomal island like <u>e</u>lements) (O’Hara et al., 2017). During the course of ICP1 infection, the integrated PLE excises from the chromosome (McKitterick and Seed, 2018), replicates (Barth et al., 2020), hijacks ICP1 machinery to package its own genome (Netter et al., 2021) and can accelerate lysis of the host *V. cholerae* host (Hays and Seed, 2020), halting spread of ICP1 to neighboring cells. Five distinct, yet highly similar PLEs have been detected in *V. cholerae* isolates going back to 1949 and over time each variant rose to prominence before being replaced by another (O’Hara et al., 2017). This succession is likely reflective of the fact that ICP1 has also evolved counter-defenses against PLEs, including an endonuclease called Odn (Barth et al., 2021) and, remarkably, a fully functional CRISPR-Cas system (Seed et al., 2013). However, the presence of these two anti-PLE elements also fluctuate over time as they target different PLEs and do not co-occur in the same genome (Barth et al., 2021). The earliest ICP1 isolates from Bangladesh in 2001 possess Odn (Seed et al., 2011), while CRISPR-Cas was first was detected in an ICP1 isolate from 2003 (Boyd et al., 2021). Between 2006 and 2017 all ICP1 isolates from Bangladesh were CRISPR(+) (McKitterick et al., 2019b). However, contemporary co-evolutionary trends of ICP1 and PLE are currently unknown.

A recent study from our lab of *V. cholerae* isolates from cholera patient stool samples in Bangladesh between 2016 and 2019 found evidence of temporal fluctuations in ICP1’s antagonism with other mobile genetic elements, SXT integrative and conjugative elements (LeGault et al., 2021b). Considering these findings, we were interested in examining how the dynamics between PLE and ICP1 have progressed during this time period. Surprisingly, we found that after dominating the epidemic landscape for nearly a decade (McKitterick et al., 2019b), PLE disappeared almost completely in 2018. This disappearance coincided with a switch from CRISPR-Cas back to *odn* among the temporally concurrent ICP1. A phylogenetic comparison of all 148 *V. cholerae* isolates during the surveillance period revealed an abrupt and nearly total shift from one *V. cholerae* lineage to a new lineage lacking PLE. Remarkably, this transition occurred over the course of only a few months. We briefly detected a novel PLE, which responds to ICP1 infection and is resistant to the anti-PLE nuclease Odn. An expanded analysis of >3000 *V. cholerae* genomes revealed the existence of additional novel PLEs and provides a clearer picture of PLE succession both geographically and temporally. We also compared *V. cholerae* isolates from within individual patients revealing heterogenous intra-patient PLE(+)/PLE(-) populations. Similarly, heterogenous ICP1 phages possessing CRISPR-Cas or Odn were isolated from a single patient and exhibited evidence of *in situ* recombination, underscoring the potential for rapid phage evolution in the human gut. These findings reinforce our understanding of the successive nature of *V. cholerae* evolution and suggest that ongoing surveillance of *V. cholerae*, ICP1 and PLE in Bangladesh is important for tracking genetic developments relevant to pandemic cholera that can occur over relatively short time scales.

## Results

### Clinical surveillance of *Vibrio cholerae* and a predatory bacteriophage uncovers regressive shifts in the arms race

We analyzed 239 stool samples from patients in Bangladesh with suspected cholera infections as determined by a rapid diagnostic test commonly referred to as a ‘dipstick’ test. These samples were recently interrogated to study the temporal dynamics of SXT integrative and conjugative elements as well as the mechanisms of phage counter-adaptation (LeGault et al., 2021b). Samples were collected between November 2016 and September 2019 with 119 originating from the capital city Dhaka and 121 from a rural city Mathbaria located on the Bay of Bengal coast (Figure 1A and Table S1).

**Figure 1.**
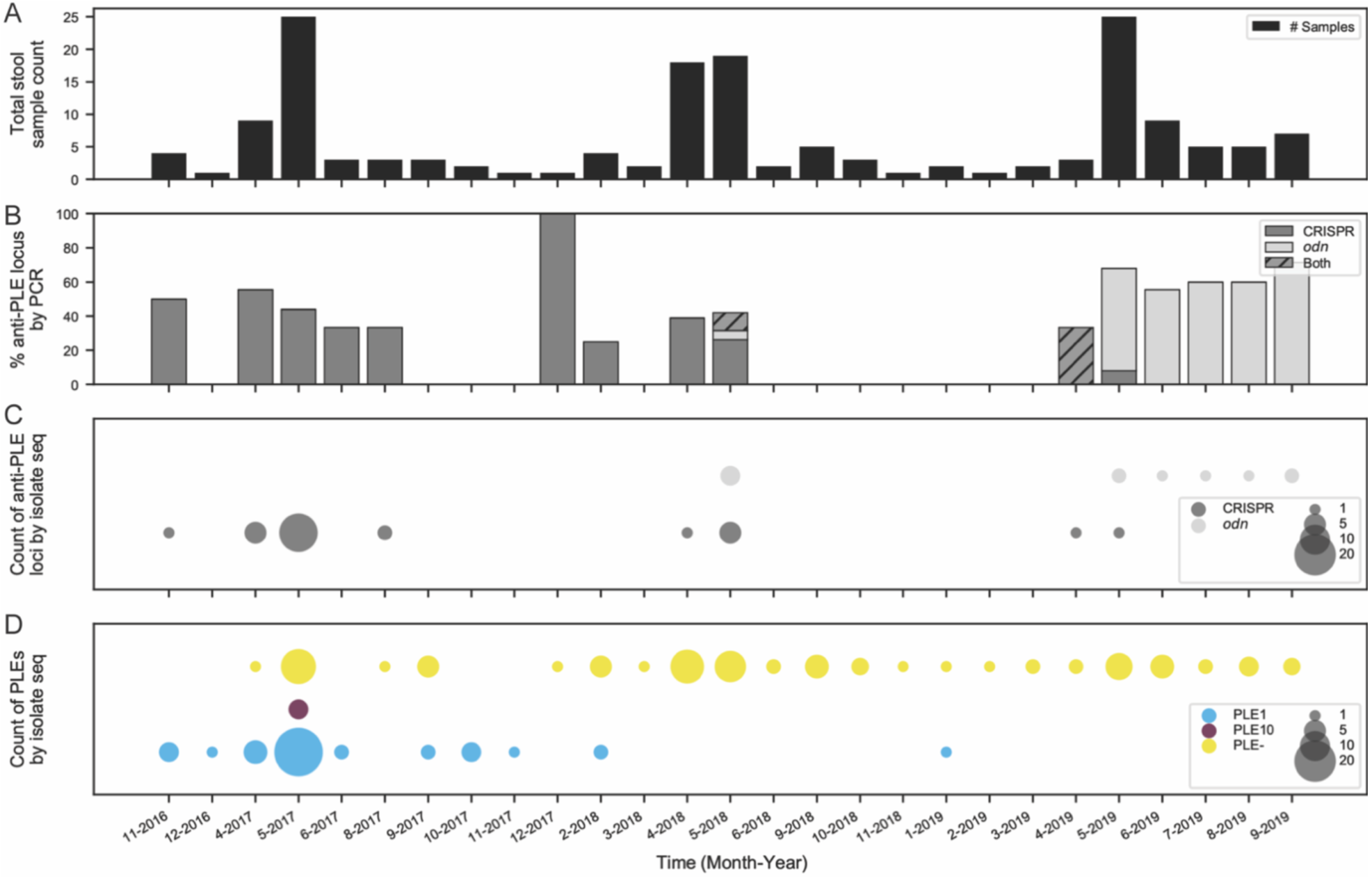
Surveillance of *V. cholerae* PLE and ICP1 anti-PLE loci in Bangladesh. A) Total counts of patient stool samples collected and analyzed by month and year of collection (239 total samples) (Table S1). B) Percentage of stool samples in a given month that screened PCR positive for ICP1-encoded anti-PLE counter-defense loci CRISPR-Cas (n=36), *odn* (n=32) or both (n=3). C) Number of whole-genome sequenced ICP1 isolates (44 total) from stool samples over the surveillance period by the type of anti-PLE counter-defense locus CRISPR-Cas or *odn*. D) Number of whole-genome sequenced *V. cholerae* isolates (148 total) from stool samples over the surveillance period by PLE variant type. PLE-indicates no PLE detected.

Interestingly, ICP1 phages, which previously encoded only CRISPR-Cas in Bangladeshi stool samples after 2006 (Angermeyer et al., 2018; McKitterick et al., 2019b), transitioned back to encoding *odn* around May 2018. Bulk metagenomic gDNA was extracted from 197 of the stool samples and used as a template for PCR to detect the presence of ICP1 by screening for *odn* and CRISPR. This screen revealed that 71 of these 197 samples were positive for ICP1: with 36 (50.7%) samples possessing CRISPR, 32 (45.1%) possessing *odn*, and three (4.2%) having a mix of both type of anti-PLE loci (Figure 1B). A sampling period over ∼11 months followed when no ICP1 was detected, then in April-May of 2019, both loci were detected again. In the final four months of surveillance (June-Sept. 2019) only *odn* was detected. To further investigate this transition from CRISPR dominance to apparent *odn* dominance, we examined whole genome sequences of 44 individual ICP1 isolates from these stool samples (Table S1). These genomes corroborated the PCR results that CRISPR-Cas dominance declined while the *odn* locus began to dominate in mid 2018 (Figure 1C).

Since both loci defend ICP1 against *V. cholerae*-encoded PLEs (Barth et al., 2021), we examined whole genome sequences of *V. cholerae* (n=148) from 110 stool samples spanning this period. From the genomes we determined whether each isolate contained PLE and if so, which of the five known PLEs was present (Figure 1D). This analysis led to two unexpected results. Firstly, PLE1, which was observed to be the dominant PLE in Bangladesh during previous surveillance (McKitterick et al., 2019b), appears to have disappeared almost entirely in favor of PLE(-) strains. Interestingly, this disappearance of PLE1 roughly correlates with the period of ICP1’s transition from CRIPSR-Cas to *odn* with only a single isolate possessing PLE1 after February 2018. And secondly, we observed that four *V. cholerae* isolates from one stool sample collected in May of 2017 possessed a novel PLE variant which we have designated PLE10. This new PLE was only observed once and was not detected in any later isolates. From these data it is difficult to say whether PLE10 provided inadequate defense against ICP1 and was selected against in the population or if increased surveillance through isolate sequencing would reveal additional instances of PLE10 in other samples.

### PLEs have evolved and diversified both globally and temporally

The discovery of a novel PLE and the surprising observation that PLEs in Bangladeshi *V. cholerae* isolates have diminished in recent years prompted us to look more broadly, both geographically and temporally, at other available *V. cholerae* genomes. Previous surveillance studies tracked the occurrence of PLEs to a limited extent (∼200 genomes) (McKitterick et al., 2019b; O’Hara et al., 2017), however this was likely insufficient to fully elucidate the flux of PLEs over time and space. To build on these analyses we constructed a database of 3,363 sequenced *V. cholerae* genomes including our new surveillance isolates and raw reads or fully assembled genomes from public repositories. This collection spans over a century (1916-2019) and includes strains collected across 25 different countries (Table S2).

Using the previously discovered five PLEs as queries (O’Hara et al., 2017), we performed BLASTn searches against each genome and identified four additional PLEs that, along with PLE10, double the total number of known PLE variants from five to 10. Abundance of these 10 PLEs has shifted over time, with PLE5 being the only PLE detected before 1987, followed by the other PLEs culminating in the more recent dominance of PLE1 (Figure 2A). Previously, PLE5 had been observed as early as 1949, but here we detected PLE5 in an isolate from 1931, increasing the known time range of PLEs circulating in *V. cholerae*. Generally speaking, the previous pattern of temporal succession, where one PLE dominates for a time before being supplanted by another PLE (McKitterick et al., 2019b; O’Hara et al., 2017) was observed in this larger collection of isolates, corroborating previous analyses. However, we observed more temporal overlap of PLEs than has previously been appreciated. Furthermore, we observed that while PLE5 had appeared to go extinct among clinical isolates in 1991, three additional PLE5 positive isolates were detected in 2016-2017 after a nearly 30-year ‘absence’, pointing to unsampled reservoirs in nature where genotypes may persist for some time.

**Figure 2.**
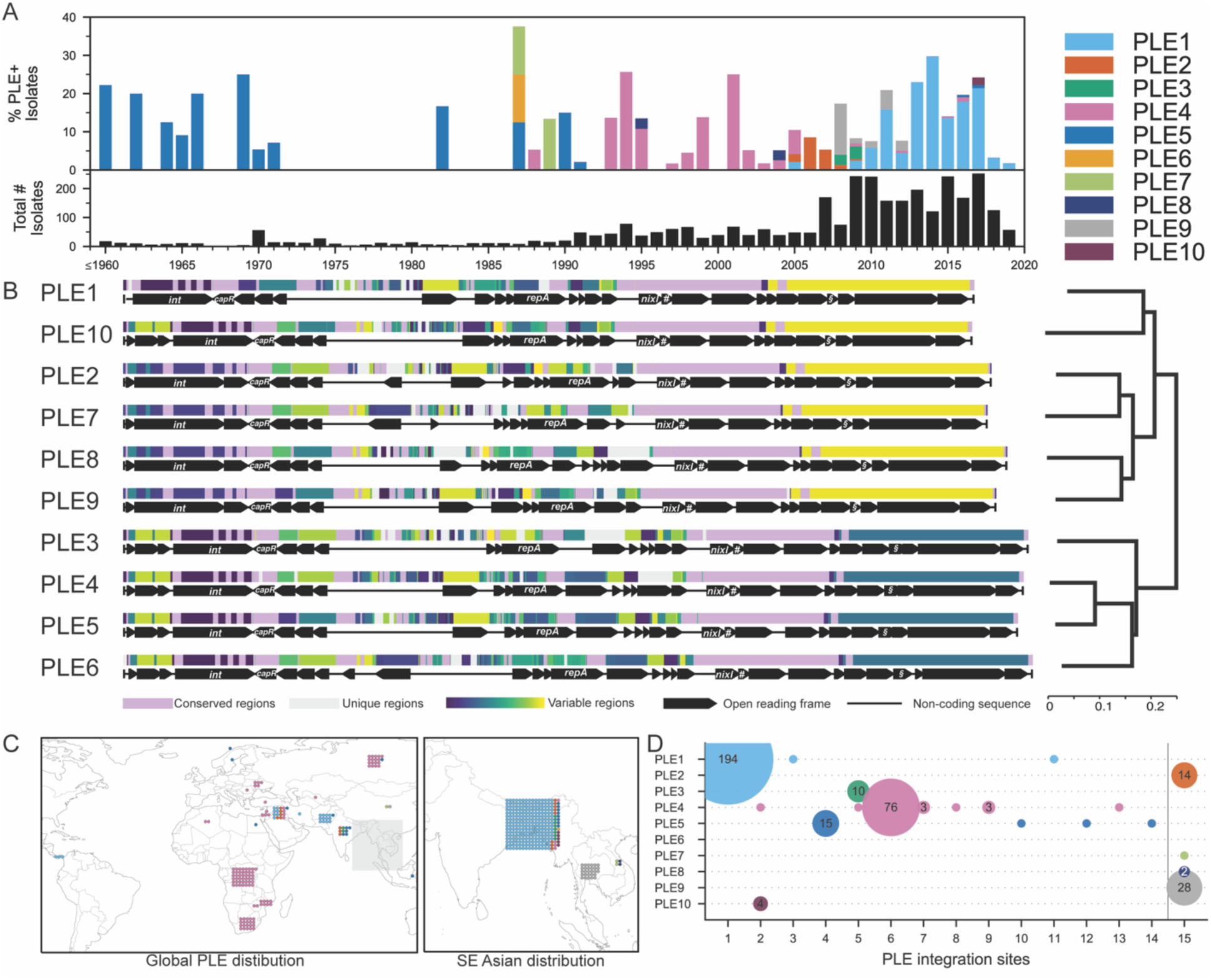
Genetic, temporal and spatial variability of PLEs in >3000 *V. cholerae* genomes collected between 1916-2019. A) Timeline of PLE-positive strains colored by PLE variant type from 3363 *V. cholerae* whole-genome sequences. The lower histogram provides the total number of strains per year, while the upper histogram represents a stacked percentage of those that are positive for each PLE. Because of the low number of sequenced genomes from isolates prior to 1960, those genomes were collapsed into a single bar. Metadata for all strains analyzed can be found in Table S2. B) Global alignment of all 10 PLEs using progressiveMauve. The mauve-colored regions are conserved across all PLE sequences, while the light grey regions are unique to that specific variant. All other regions vary in conservation between two or more PLEs and were assigned random colors from the variable palette. Annotated genes are shown to scale as black arrows, several genes with known functions are labeled, #=*stiX* and §=*lidI*. The alignment’s final guide tree on the right provides an approximation of phylogenetic distance and determines the order of sequences. The scale represents the number of nucleotide substitutions per base pair. C) Geographic locations by total number of PLE+ isolates on a global scale (left) excluding Southeast (SE) Asia in grey box and zoomed into that region (right). The color of the dot indicates the PLE variant as in panel (A).

Some of the newly discovered PLE variants, however, only appeared for a short time, such as PLE6 which was detected once in Bangladesh in 1987, and PLE8 in Vietnam in 1995 and 2004. We also detected one possible phage satellite with partial genetic relationship to PLEs, as has been described in non-cholera Vibrios recently (LeGault et al., 2021a), however its sequence could not be fully resolved from available data. It is highly likely that we still have not captured the full extent of PLE diversity and variant flux over time as *V. cholerae* isolate sampling was sporadic and limited until recent decades (with over half of all isolates collected in only the last nine years). For example, in this current surveillance effort we observed PLE10 in only a single patient sample which likely would not have been observed at all with slightly less sampling depth. Whether these short-lived PLEs (and divergent satellites) are somehow less adept at parasitizing ICP1, are selected against by specific anti-PLE counter-defenses, or are victims of simple stochastic fluctuations, is unknown.

Additionally, transmission bottlenecks that likely occur in the course of regional outbreaks may constrain PLEs geographically (Figure 2C), such as in the two most recent outbreaks in Haiti (Piarroux et al., 2011) and Yemen (Weill et al., 2019) where no PLE was detected. The most notable exception being Bangladesh where seven of the 10 PLE variants have been detected. While this may be partially due to the sampling focus on this region as a significant fraction (∼20%) of the *V. cholerae* genomes analyzed were from there, it may also be due to its juxtaposition with the Bay of Bengal where the majority of *V. cholerae’s* diversification and evolution is predicted to occur.

Having found additional PLEs, we then performed a whole genome alignment of all 10 PLE variants which was used to generate a phylogenetic tree (Figure 2B). All PLEs are broadly syntenic and share numerous regions of nucleotide conservation. Annotation and sequence comparison of open reading frames confirmed that the newly detected PLEs 6-10 possess genes key to PLE function that have been characterized in the previously studied PLEs 1-5. These include *nixI*, which encodes a nickase that interferes with ICP1’s ability to replicate and a supplementary gene *stiX* which increases NixI activity (LeGault et al., 2021a), *lidI*, which can accelerate lysis of the *V. cholerae* host cell (Hays and Seed, 2020), and *capR*, which acts to repress ICP1’s capsid morphogenesis operon (Netter et al., 2021). The conservation of these genes indicates that the newly identified PLEs play similar roles as PLEs 1-5 as parasites of ICP1. Two additional genes indicate that these PLEs are also still functional as mobilizable genetic elements: an integrase that catalyzes PLE excision from the host genome and integration following transduction, and a replication initiation factor that controls PLE replication.

PLEs encode one of two integrases (Figure S1), the PLE1-type (found in PLEs 1,3,4,5,6,10) which directs integration into a *Vibrio cholerae* repeat (VCR) sequence in the superintegron and recognizes ICP1-encoded PexA as a recombination directionality factor to catalyze PLE excision (McKitterick and Seed, 2018), and the PLE2-type (found in PLEs 2,7,8,9) integrates into an M48 family metallopeptidase gene (*vca0581*) and does not interact with PexA. VCRs are ∼124bp repeat elements that are interspersed between gene cassettes within the superintegron (Barker et al., 1994). VCRs are extremely conserved, which could allow PLEs with the PLE1-type integrase to integrate into any single VCR following horizonal transfer resulting in isogenic PLEs flanked by different genes in the superintegron. Previous analyses show that PLE1 integrates into various distinct VCRs under laboratory conditions, but such variability has not been observed among PLE(+) *V. cholerae* isolates from clinical specimens, suggesting that PLE transmission is largely vertical in nature (McKitterick et al., 2019b; O’Hara et al., 2017). To determine if this finding held true, we extracted the genomic regions flanking PLE for every PLE positive *V. cholerae* isolate in our dataset. While no flanking differences were observed among the PLEs with PLE2-type integrases, we identified 14 unique VCR integration sites for the PLE1-type integrase positive *V. cholerae* isolates in our database (Figure 2D and Table S3). And while most PLE variants appeared to have a single integration site, either through preferential horizontal integration or vertical transmission, we did detect multiple sites for PLE1, PLE4, and PLE5 indicating likely horizontal acquisition events in nature.

The PLE-encoded replication initiation protein RepA has a conserved C-terminus across all PLEs, but varies at the N-terminus which directs DNA binding to the PLE origin of replication (ori) (Barth et al., 2020). Previous work found that RepA’s N-terminus for PLEs 1-5 fell into two groups, one with PLEs 1,4,5 and the second with PLEs 2&3. Diversification of the PLE replication module (comprised of a compatible RepA and ori) is hypothesized to be driven by ICP1’s nuclease Odn which mimics the PLE1,4,5-type RepA protein to bind to the cognate ori and then cleave it. For PLE to escape Odn by modifying its ori, it must possess a compatible RepA N-terminus as well as the conserved C-terminus which is hypothesized to facilitate recruitment of the replisome machinery from ICP1 (Barth et al., 2020). Including the five new PLEs described here, phylogenetic analysis revealed the existence of a novel third RepA N-terminal variant found in PLEs 6,8,9,10 (Figure S2). In support of the designation of a third type of PLE replication module, previously validated ori sequences in PLE1-5 were not observed in PLEs with the PLE10-type RepA protein, indicating that these PLEs harbor a compatible unique ori sequence that would be resistant to Odn-mediated cleavage.

### PLE10 responds to ICP1 infection and is resistant to Odn

All previously studied PLEs (PLEs 1-5) are triggered by ICP1 infection to excise, replicate and package themselves into transducing particles using ICP1’s machinery (Figure 3A) (O’Hara et al., 2017). Based on genetic similarity to PLEs 1-5, we hypothesize that the five new PLEs described here respond similarly to ICP1 infection. Unfortunately, of the new PLE variants we only had access to *V. cholerae* strains possessing PLE10 and therefore experimentally assessed how this variant responds to ICP1 infection.

**Figure 3.**
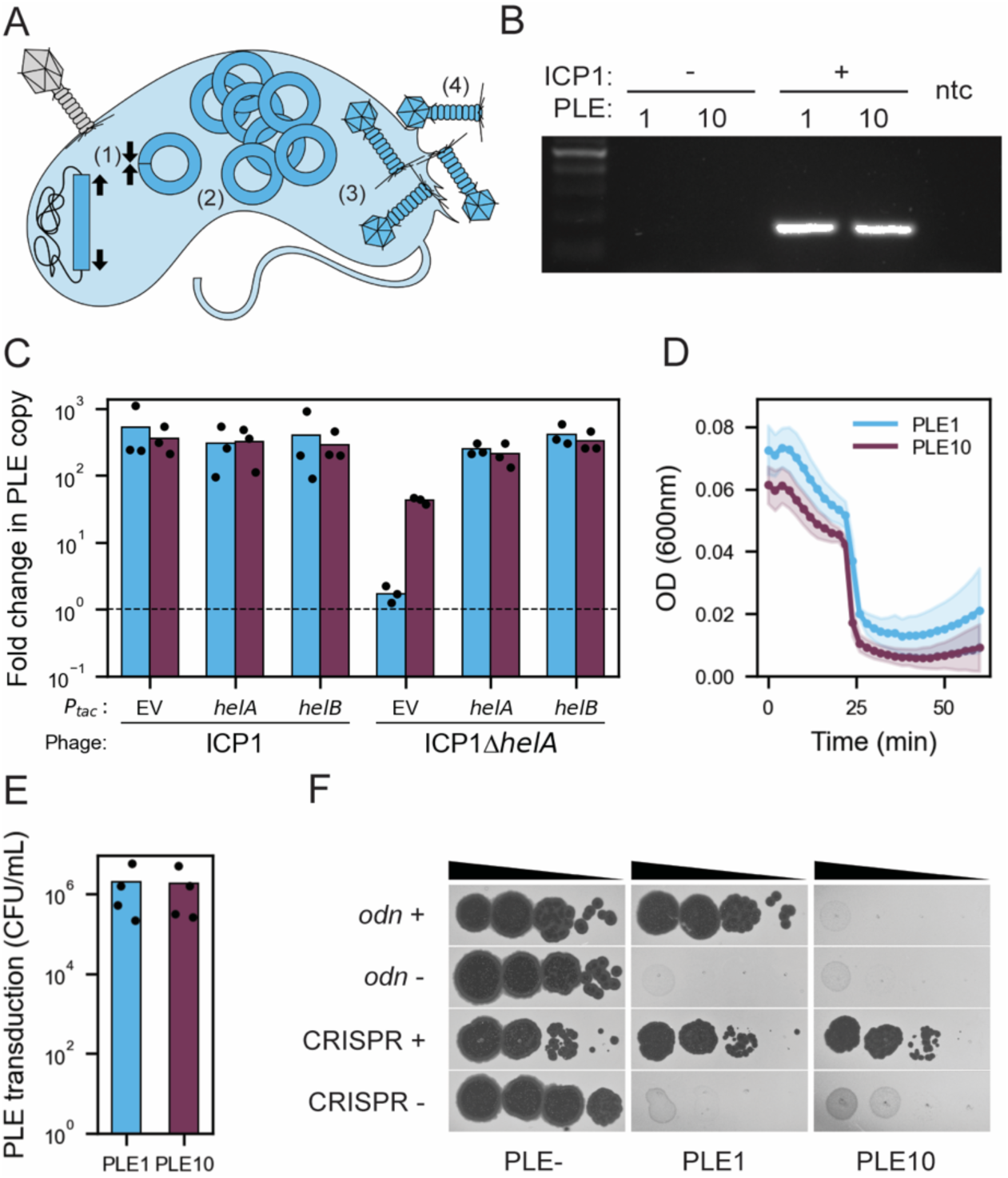
PLE10 exhibits excision, replication and packaging in response to ICP1 infection and evades the anti-PLE nuclease Odn. A) Model showing the steps of the PLE-ICP1 interaction in a *V. cholerae* host cell. (1) The integrated PLE excises and circularizes upon ICP1 infection. (2) PLE replicates and (3) is packaged into modified phage particles. (4) PLE transducing-particles are released through cell lysis which occurs on an accelerated timeline. B) Agarose gel of PCR products to detect circularized PLE in uninfected *V. cholerae* and following infection by ICP1^2006^ lacking CRISPR-Cas (the phage isolate previously used to probe PLE circularization (McKitterick and Seed, 2018)). C) Quantification of change in PLE1 and PLE10 copy number 30 minutes after infection with ICP1^2006^ lacking CRISPR-Cas or the Δ*helA* mutant (the phage isolate previously used to probe PLE replication (Barth et al., 2020; O’Hara et al., 2017))). IPTG inducible plasmid constructs (P_*tac*_) were induced prior to phage infection. EV is the empty vector control. The dashed line indicates no change in copy number. D) Lysis of *V. cholerae* harboring PLE1 or PLE10 following infection by ICP1^2006^ lacking CRISPR-Cas (the phage isolate previously used to probe lysis kinetics (Barth et al., 2020; O’Hara et al., 2017). E) PLE transducing-particles generated during infection with ICP1^2006^ lacking CRISPR-Cas (the phage isolate previously used to probe PLE transduction (O’Hara et al., 2017)).

To determine if this novel PLE10 variant indeed functions in a similar way to known PLEs, we performed several assays in tandem with PLE1. First, a circularization PCR showed that PLE10 excises and circularizes during ICP1 infection (Figure 3B) as was expected due to the similarity of the PLE1 and PLE10 integrases (Figure S1). Second, we observed that PLE10 replicates in response to ICP1 infection (Figure 3C). This confirms that the divergent PLE10 replication module is functional and similarly exploits ICP1 machinery to promote PLE replication. Interestingly, although PLEs 1-5 do not replicate during infection by ICP1 mutants lacking an accessory SF1B-type helicase (*helA* or *helB*) (McKitterick et al., 2019a), PLE10 does replicate to low levels in the absence of this ICP1-encoded helicase (Figure 3C), suggesting that the divergent replication module allows for additional flexibility in PLE’s reliance on ICP1’s replication machinery. Third, we observed that PLE10, which possesses a *lidI* homologue (Hays and Seed, 2020) also exhibited an accelerated lysis phenotype like PLE1 following ICP1 infection (Figure 3D). And fourth, PLE10 mobilized and transduced its genome to naive *V. cholerae* hosts during infection at rates similar to PLE1 (Figure 3E). These results demonstrate that PLE10 exhibits the anticipated response to ICP1 infection, namely excision, replication and transduction.

As ICP1 is known to possess two anti-PLE counter-defense loci (*odn* and CRISPR-Cas), we tested the ability of ICP1 with either loci to plaque on *V. cholerae* having either PLE1 or PLE10. PLE1 is susceptible to ICP1 possessing CRISPR-Cas (those with a spacer specific to PLE1) (Seed et al., 2013), and one hypothesis is that this new PLE10 variant may have had an advantage over PLE1 through the acquisition of an anti-CRISPR, although none have been found previously in PLEs. However, we found that the CRISPR-Cas positive phage, isolated from the same stool sample as the PLE10 *V. cholerae* isolates and naturally possessing spacers specific to both PLE1 and PLE10 (Table S4), was able to overcome PLE10 during infection and that this activity was dependent on ICP1’s CRISPR-Cas system (Figure 3F). This was an identical result to the PLE1 control, which we expected to be overcome by the same CRIPSR-Cas(+) phage. The inability to block plaque formation by CRIPSR-Cas(+) phage indicates that PLE10 has no anti-CRISPR protein that might provide an evolutionary advantage over PLE1.

In contrast, an ICP1 phage with *odn* was unable to form plaques on a PLE10(+) host, but was able to form plaques on an otherwise isogenic host harboring PLE1 (Figure 3F). Purified Odn also did not exhibit nucleolytic activity against a probe amplified from the ORF-less region of PLE10, compared to a probe from PLE1 which was cleaved as expected (Figure S3). Collectively these data are consistent with the specificity of the origin-targeting behavior of Odn (Barth et al., 2021) and the divergent PLE10 replication module conferring protection from Odn-mediated cleavage. The *odn*(+) phage tested for its capacity to form plaques on PLE10 was from a more recent 2019 stool sample, a time period in which *odn*-positive ICP1 phages were dominant in our surveillance (Figure 1C). However, despite the observation that PLE10 is resistant to this *odn*+ phage, it did not have an apparent selective advantage over PLE1 during this surveillance period (Figure 1D).

### Colony and plaque screens reveal intra-patient heterogeneity

Preliminary PCR screening of metagenomic DNA from stool samples revealed some samples with a positive signal for PLE, but from which we later recovered *V. cholerae* isolates that were PLE(-). This finding prompted us to interrogate several samples further to better understand the ratios of PLE(+) to PLE(-) strains within intra-patient populations. We selected three samples from May 2017 (D19, D28, D35) and one from September 2017 (D55), as both timepoints also possessed a mix of PLE+ and PLE-isolates between patient samples (Figure 1D). We performed PCR on a large number of colonies (n=60-130) picked from each of these samples and observed that all samples were indeed heterogenous (ranging from 6-97% PLE(+)) (Figure 4A). We additionally chose one sample from May 2017 from which all (n=4) sequenced isolates were PLE(+) to see if PLE heterogeneity could be caused by stochastic loss under laboratory conditions. PCR of 218 colonies revealed all to be PLE(+) (Figure 4A), indicating that PLE loss was highly unlikely to occur as a result of the outgrowth and isolation procedures we used. These results reinforce previous discoveries that *V. cholerae* can diversify to some extent during human infection (Levade et al., 2017), although whether this finding indicates a loss or gain of PLE is unclear.

**Figure 4.**
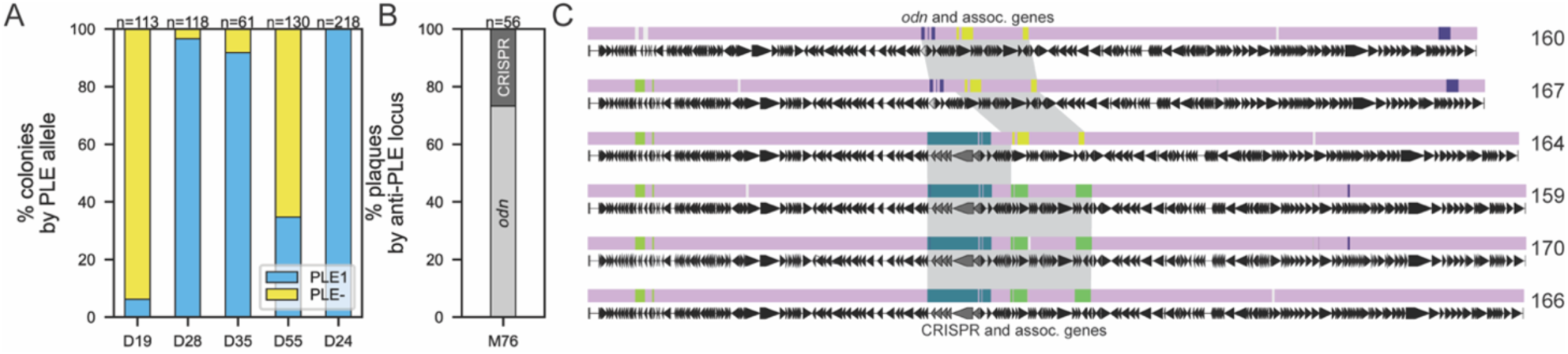
PLE(+)/PLE(-) *V. cholerae* and ICP1 anti-PLE loci heterogeneity within patients. A) The fraction of total colonies with and without PLE for *V. cholerae* isolates within individual stool samples as determined by PCR. The number of colonies screened from each patient sample (given an arbitrary number by date of isolation with the prefix D are from icddr,b in Dhaka) are given above each bar. B) The fraction of ICP1 plaques with the anti-PLE locus indicated recovered from one stool sample from Mathbaria (M76) as determined by PCR. C) Alignment of six ICP1 isolates from stool sample M76 (panel B) using progressiveMauve. The mauve-colored regions are conserved across all ICP1 sequences, while the light grey regions are unique to that specific isolate. All other regions vary in conservation between two or more isolates and were assigned random colors from the variable palette. Annotated genes are shown as black arrows, Cas and *odn* genes are shown in grey. The grey shading highlights regions with *odn* with associated genes (in phages 160 and 167) and CRISPR-Cas with associated genes (in phages 159,170 and 166) at ≥95% sequence identity. Isolate 164 demonstrates recombination between the CRISPR-Cas region and the *odn*-associated region. See Table 1 for complete description of phage isolates.

Similarly, we performed PCR on a large number of ICP1 plaques from sample M76 (May 2018) in which the initial screen of metagenomic DNA extracted from stool revealed a mixed signal of both ICP1-encoded *odn* and CRISPR-Cas anti-PLE loci. This was also the first instance of *odn* reappearance (Figure 1B) in this surveillance period. All previously discovered ICP1 isolates have either *odn* or CRISPR-Cas, but never both (Angermeyer et al., 2018; Barth et al., 2021), and these counter-defense mechanisms have always been in the same location within the genome. It is an intriguing possibility that some rare isolates could carry both *odn* and CRISPR-Cas, but PCR results from purified plaques (n=56) from this sample showed that members of this intra-patient ICP1 population indeed only had CRISPR-Cas or *odn* (Figure 4B).

**TABLE 1.**
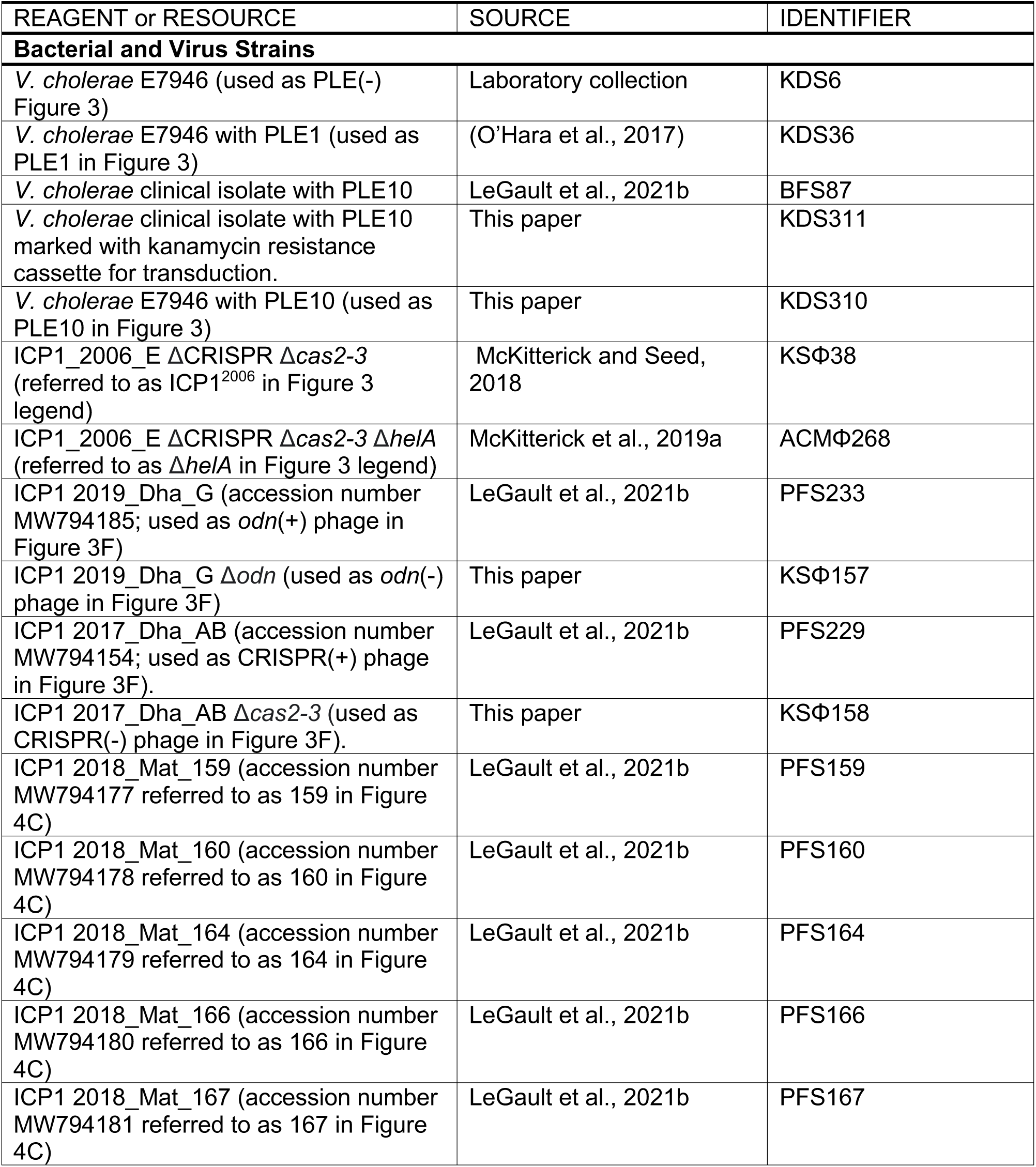
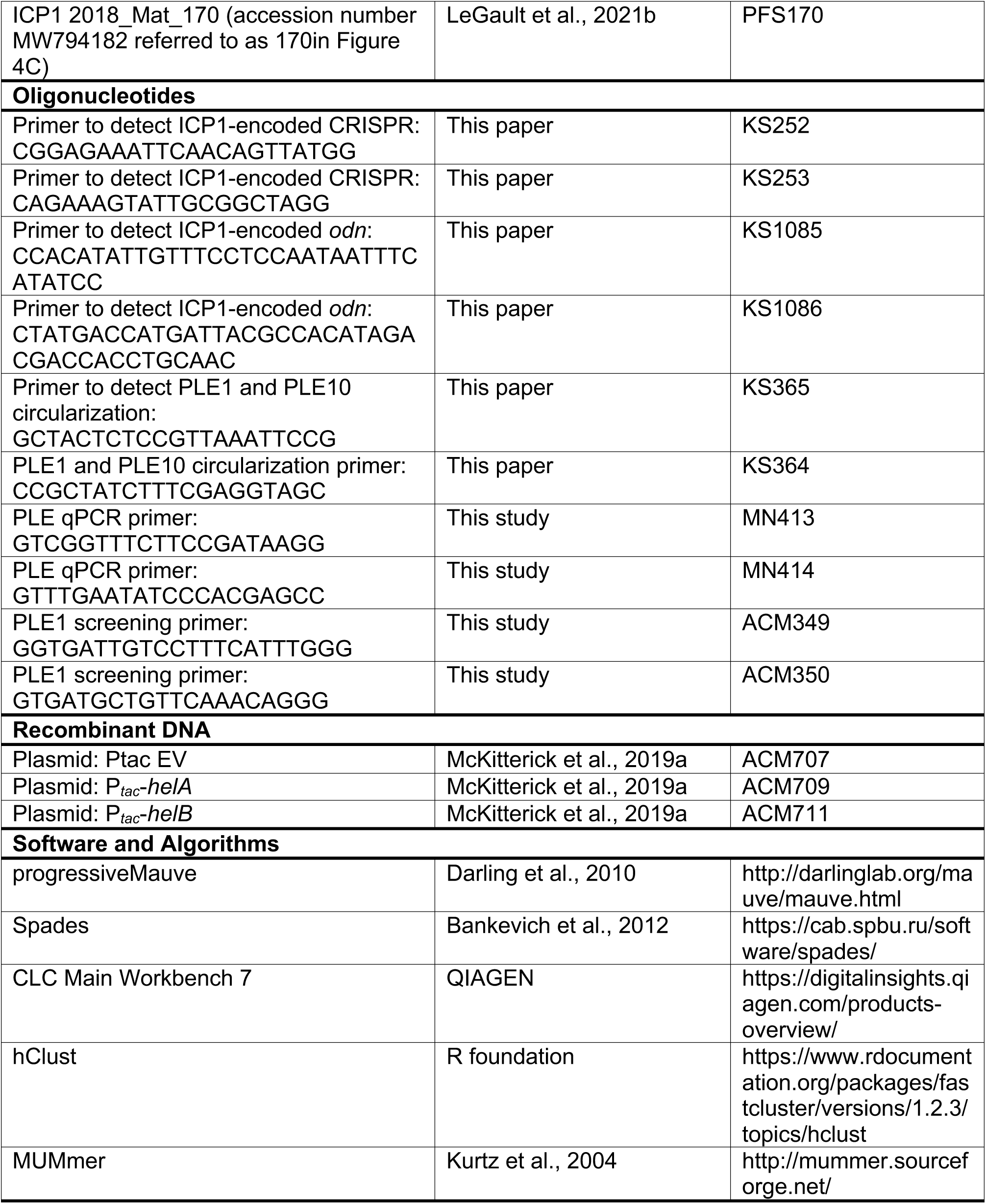

In addition to the 56 screened plaques, we also sequenced the genomes of six isolates of ICP1 from this patient sample, two *odn*(+) isolates and four CRISPR(+) isolates. Whole genome sequencing revealed that both *odn*(+) phage had a canonical downstream region typically associated with *odn* and three of the CRISPR(+) phage had the expected distinct conserved set of associated genes. However, one isolate (2018_Mat_164) harboring the CRISPR-Cas anti-PLE locus possessed the downstream region typically associated with *odn* (Figure 4C). This appears to be an example of within-patient recombination between two ICP1 genotypes and is strong additional evidence that co-infection is a likely driver of ICP1 evolution as has been demonstrated previously under laboratory conditions (Hays and Seed, 2020). To the best of our knowledge, this is the first attempt to document intra-patient diversity of vibriophages and as such the rates of *in-situ* recombination are unknown. Nonetheless, these data highlight that ICP1 evolution occurs in the human gut, giving rise to distinct ICP1 genotypes with distinctive host-ranges (Barth et al., 2021) that could differentially impact the fitness of co-circulating *V. cholerae*.

### *V. cholerae* isolates from 2016-2019 in Bangladesh sort into two distinct phylogenetic lineages

The transmission and evolution of *V. cholerae* is characterized by successive waves of clonally related strains that are replaced over time (Weill et al., 2017) and the phylogenetic relationships between these waves have generally been determined by differences in single point mutations. We observed that PLE disappeared from *V. cholerae* isolates during the course of our surveillance (Figure 1D), however we did not know if this is the result of PLE being lost from a clonal population or if the PLE(+) lineage was replaced by a distinct PLE(-) lineage. To address this question, we identified 532 sites with single nucleotide variations (SNVs) (Table S5) common across all 148 isolates (Table S6) and used this information to construct a core genome phylogenetic tree (Figure 5).

**Figure 5.**
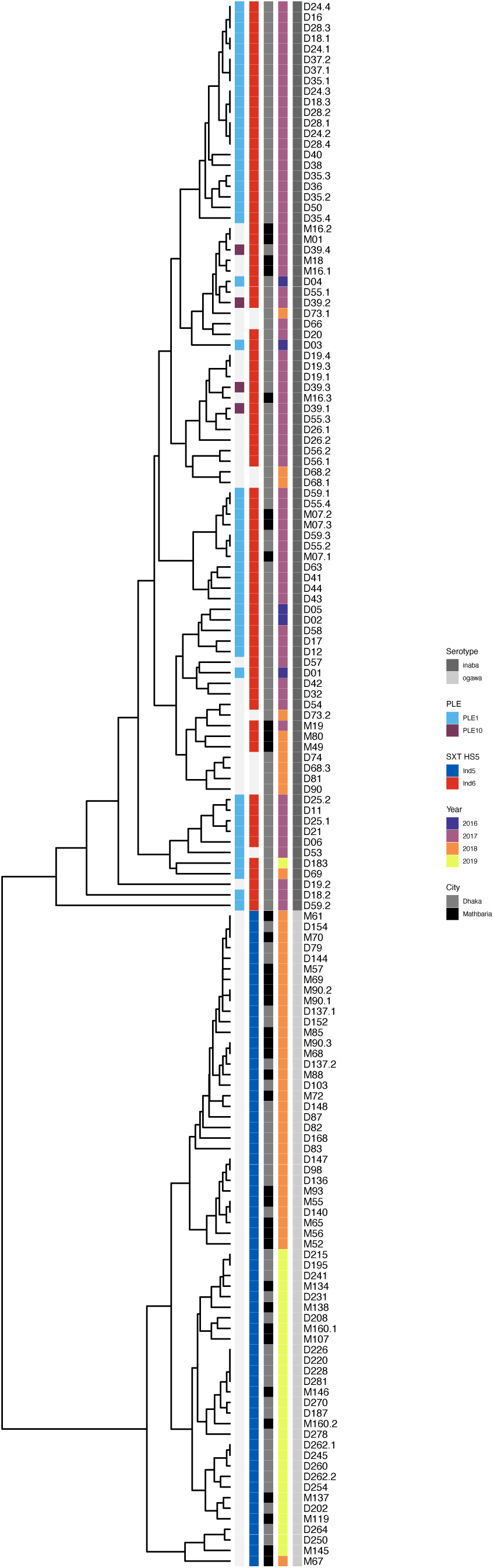
Phylogeny of *V cholerae* isolates during the surveillance period in Bangladesh reveals the replacement of a previously prevalent PLE(+) lineage with a PLE(-) lineage in 2018-2019. A cladogram of 148 *V. cholerae* isolates based on 532 variable nucleotide genomic positions (Table S5). Columns next to the tree contain metadata for (left to right) the PLE variant present, the SXT ICE present, city of origin, year isolated and serotype (based on whole-genome sequencing and analysis of *wbeT*). Strains lacking the MGE (PLE or SXT ICE) are indicated with a light grey shade. Each isolate is labeled according to the arbitrary designation of the patient stool sample it was isolated from. Samples were given an arbitrary number by date of isolation, those with the prefix D are from icddr,b in Dhaka and those with the prefix M are from Mathbaria. Decimal values indicate that multiple isolates were collected from that single stool sample (Table S6).

It was immediately clear that the *V. cholerae* isolates cluster into two highly distinct phylogenetic groups which largely coincide with the presence of PLE, suggesting that we captured a replacement event where the PLE(-) lineage emerged to dominate as opposed to witnessing the simple loss of this MGE in an otherwise persistent lineage. This was further reinforced by a previous observation in this surveillance dataset that another MGE which plays a role in phage defense, integrative and conjugative elements (ICEs) of the SXT/R391 family, exhibited a transition from a mix of *Vch*Ind6 and SXT(-), to *Vch*Ind5 over the same time period (LeGault et al., 2021b). Additionally, sequences of the *wbeT* gene, which is a part of the biosynthetic pathway for the O1 antigen, also revealed a switch from the Inaba (nonfunctional *wbeT*) to Ogawa (functional *wbeT*) serotype that perfectly tracked with the lineage replacement. In the earlier Inaba lineage, *wbeT* was disrupted by a transposon with an identical sequence and integration site across all isolates. Remarkably, this replacement sweep appears to have occurred over months instead of years with only a few isolates from the earlier PLE(+)/*Vch*Ind6(+)/Inaba lineage persisting into 2018 and only a single one in 2019. This may indicate that the later PLE(-)/*Vch*Ind5(+)/Ogawa lineage has some competitive advantage over the earlier lineage driving its sudden and dominating emergence. Also, we saw no patterns separating the sampling locations of Dhaka and Mathbaria, possibly indicating that the replacement sweep occurred rapidly geographically as well as temporally.

Finally, in addition to the samples examined by colony PCR (Figure 4A), multiple isolates were sequenced from several other samples when recoverability permitted. And while significantly more isolate sequencing would be required to draw meaningful conclusions about the full extent of intra-patient diversity, we did observe both SNV and MGE variability between isolates in at least two patient samples (D19 and D55). However, in no sample did we detect isolates from both lineages among the intra-patient isolates, perhaps suggesting that although rapid, lineage sweeps are unlikely to occur in the context of a single patient’s infection.

## Discussion

The trajectories of global cholera outbreaks are generally defined by successive waves of *V. cholerae* in which a new lineage replaces the previous lineage (Weill et al., 2017). And while it is unclear exactly what allows successive lineages to outcompete their predecessors, it is broadly theorized that each new lineage in the current 7^th^ pandemic has originated from the areas surrounding the Bay of Bengal (Mutreja et al., 2011). Therefore, developing a better understanding of the genetic variability of *V. cholerae* in this region is likely key to unlocking the mystery surrounding the cycle of global selective sweeps. Furthermore, predatory bacteriophages like ICP1 and anti-phage defenses such as PLE may play an important role in determining which lineages become pathogenically relevant in human populations. In this study we describe findings from continued surveillance of *V. cholerae* in Bangladesh and build upon previous work to further reinforce the importance of monitoring these microbial populations in communities proximal to the Bay of Bengal.

The antagonism between ICP1 and PLE results in an ongoing arms race in which ICP1 evolved means to target PLE and inhibit PLE’s anti-ICP1 activity. The earliest known ICP1 isolates have the Odn nuclease which cleaves PLE’s origin of replication (Barth et al., 2021) while PLEs likely diversified to escape Odn targeting. It was assumed, however, that the extreme flexibility and rapid adaptation provided by the later acquisition of a novel CRISPR-Cas system had given ICP1 the upper hand in this battle (McKitterick et al., 2019b). Or at least that CRISPR-Cas would be so beneficial so as to remain fixed in the ICP1 population. Remarkably though, we observed that despite the assumed superiority of a CRISPR-Cas counter-defense mechanism, a reversion occurred in the ICP1 population back to carrying the *odn* locus instead of CRISPR-Cas. During this same timeframe before reversion to Odn we also found that nearly all *V. cholerae* isolates lacked PLE. Whether or not these two nearly simultaneous changes are functionally connected is unclear. Perhaps the much larger CRISPR-Cas locus is too burdensome for the phage to encode when PLE no longer threatens its viability, or maybe *odn* is more effective at fully restricting PLE activity for the PLEs that it can target, or possibly both (Barth et al., 2021).These hypotheses suggest that PLE was lost from the *V. cholerae* population first and in the absence of that selective pressure, ICP1 quickly reverted to a more favorable genotype.

A similar ‘reversion’ event was previously observed in these same surveillance samples in which the promoter for a gene (*orbA*) that inhibits anti-phage components of the SXT ICE *Vch*Ind5 was missing in ICP1 isolates while *Vch*Ind5 was absent from the *V. cholerae* population and a different SXT ICE dominated. However, when *Vch*Ind5 became dominant in the population around mid 2018 (Figure 5), phage isolates with *orbA* and its functional wild-type promoter also re-emerged as the dominant ICP1 genotype (LeGault et al., 2021b). It is possible that ICP1’s robust accessory genome allows it to keep pace with emergent *V. cholerae* lineages, and reemergence of historical genotypes suggests unsampled reservoirs of ICP1 diversity much like we expect for *V. cholerae*. And if future lineage shifts cause PLE to re-emerge in the *V. cholerae* population, it may also mean that CRISPR-Cas, or perhaps yet another anti-PLE mechanism, will again supplant Odn in the ICP1 population. However, this hypothesis is potentially weakened by the discovery of a new PLE in one patient stool sample approximately a year before the shift in ICP1’s anti-PLE loci. The new PLE (PLE10) was shown to be functional as a mobile element in response to ICP1 infection but was also resistant to *odn* targeting by having a novel origin of replication. In this case, why then did PLE10 not proliferate in the *V. cholerae* population after ICP1 had switched to *odn*? It is impossible to say for certain, but it may have some unknown competitive disadvantage or was simply unable to spread through the population while CRISPR-Cas with PLE10 spacers was still co-occurring. It is further possible that the pressures inherent in the dynamics of ICP1-PLE interactions are not responsible for driving *V. cholerae* lineage selection and PLE10 was simply not present in the lineage that swept to dominance in 2018.

In further investigating the genomics of this lineage switch among the *V. cholerae* surveillance isolates we observed two very distinct clades based on SNVs. And while the SNV analysis didn’t reveal any candidate mutations that would indicate an obvious competitive advantage of the new lineage, a sizable number of the non-synonymous variations were perfectly separated between the two lineages (Table S7). This may indicate that the lineage shift reflects an emergence of a distinct *V. cholerae* variant that already had these mutations in its genome. Additionally, the serotype switch from Inaba to Ogawa also tracked precisely with the two lineages. This tendency of *V. cholerae* lineages to change serotype between Inaba and Ogawa, or vice versa, has been well-documented. This is in large part due to the ease in which serotype agglutination tests can be performed, and has revealed these shifts to be very common (Koelle et al., 2006). It has been documented globally (M. T. Alam et al., 2016; Dorman et al., 2020) as well as in Bangladesh for almost five decades (Longini et al., 2002), including a time range that overlaps in part with our surveillance period and agrees with the observation made here (Baddam et al., 2020). Despite this long history, it is still unknown what drives the Inaba/Ogawa serotype switching in *V. cholerae*. As the function of the *wbeT* gene is to modify the O1-anitgen, it has been hypothesized that phage predation could be a selective pressure (Karlsson et al., 2016), however ICP1 (which uses the O1 antigen as its receptor) is insensitive to this modification (Seed et al., 2011) and there is no evidence to indicate that the other two known vibriophages frequently isolated from patient stool samples (ICP2 and ICP3) are impacted by such a modification. An alternative hypothesis is that O1 antibodies created as an immune response to *V. cholerae* in patient populations could select against whichever serogroup is currently circulating. While this has not been conclusively tested, a recent study found that patients challenged with an Inaba strain developed antibodies that cross-reacted effectively with the Ogawa serotype (Hossain et al., 2019), suggesting a limited selective advantage to serotype switching in evading human immunity.

We did find some evidence of possible diversification within patients by detecting mixed populations of PLE(+) and PLE(-) colonies from several patient samples as well as in sequenced isolates from one patient that varied by several SNVs. These observations are reinforced by previous discoveries that found genetic variation during human infection (Levade et al., 2017) and even the emergence of hypermutator phenotypes (Levade et al., 2021). Whether this indicates an *in situ* loss/gain of PLE or co-infection with distinct strains is unknown. We also found evidence of bacteriophage heterogeneity leading to recombination within a single patient. We are not aware of this being observed in nature before, and there is experimental evidence of ICP1 recombination under laboratory conditions (Hays and Seed, 2020). The evolutionary variation in the *V. cholerae*/ICP1/PLE system is likely apparent, and perhaps more so, in the aquatic reservoir. A deeper investigation of individual patient samples as well as longitudinal surveillance and environmental samples will be important to begin to fully understand the dynamics at play in driving *V. cholerae* evolution and lineage selection.

## Acknowledgments

We are especially thankful to icddr,b hospital and lab staff for support. We thank members of the Seed lab for critical feedback and thoughtful discussion regarding this manuscript. Funding: The project described was supported by Grant Numbers R01AI127652 and R01AI153303 (K.D.S.) from the National Institute of Allergy and Infectious Diseases and its contents are solely the responsibility of the authors and do not necessarily represent the official views of the National Institute of Allergy and Infectious Diseases or NIH. K.D.S. is a Chan Zuckerberg Biohub Investigator and holds an Investigators in the Pathogenesis of Infectious Disease Award from the Burroughs Wellcome Fund. icddr,b gratefully acknowledges the following donors, which provide unrestricted support: Government of the People’s Republic of Bangladesh, Global Affairs Canada, Swedish International Development Cooperation Agency (SIDA), and the Department for International Development, UK Aid.

## Author contributions

A.A., S.G.H. and K.D.S. designed the study. A.A. wrote the manuscript which was edited by K.D.S. A.A. performed the sequence and bioinformatic analyses. S.G.H. performed PCR screens. S.G.H., M.H.T. and K.D.S performed experiments to characterize PLE activity. F.J., M.S., and M.A. provided essential Resources. M.A. and K.D.S acquired funding. All authors reviewed and edited the manuscript.

## Materials & Methods

### Bacterial Growth Conditions

The bacterial strains and plasmids used in this study are listed in Table 1. All bacterial strains were grown at 37°C in LB (Fisher) with aeration or on LB agar plates. Antibiotics were used when appropriate (100 µg/mL streptomycin, 75 µg/mL kanamycin; 2.5 µg/mL chloramphenicol for *V. cholerae* and 25 µg/mL for *E. coli*). Ectopic expression constructs in *V. cholerae* were induced 20 minutes prior to ICP1 infection with 1 mM Isopropyl β-D-1-thiogalactopyranoside (IPTG) and 1.5 mM theophylline.

### Phage Growth Conditions

The phages in this study are listed in Table 1. Bacteriophages were propagated on *V. cholerae* and harvested via polyethylene glycol precipitation (Clokie et al., 2009) or media exchange on Millipore’s Amicon Ultra centrifugal filters (Bonilla et al., 2016) and quantified via the soft agar overlay method (Clokie et al., 2009).

### Isolation of Bacteria, Phage and Total DNA from Stool

The collection of stool samples and isolation of *V. cholerae* and phages interrogated in this study was recently described (LeGault et al., 2021b). Briefly, de-identified rice water stool (RWS) from patients presenting with diarrheal disease were tested for the presence of *V. cholerae* via Crystal VC Rapid Dipstick test (RDT; Span Diagnostics, Surat, India) at icddr,b Dhaka hospital and Government Health Complex of Mathbaria, Pirojpur. RDT-positive stool samples were mixed with glycerol and frozen for transport to University of California, Berkeley. *V. cholerae* was isolated from RDT-positive stool samples onsite in Bangladesh following enrichment in alkaline peptone water (APW) and plating on taurocholate tellurite gelatin agar (TTGA) (Difco). Additional isolates of *V. cholerae* from these RWS samples were obtained following APW outgrowths and culturing thiosulfate-citrate-bile salts-sucrose agar (Fisher), and Vibrio ChromoSelect Agar (Sigma)) at the University of California, Berkeley. To isolate phages, dilutions of RWS and APW outgrowths were mixed with log-phase *V. cholerae* and plated in 0.3-0.5% LB top agar. Total DNA was extracted from stool samples using the DNeasy PowerSoil kit (Qiagen) using 333 μL of stool following a modified extraction protocol. Briefly, 200 μL of bead solution was removed and replaced with 200 μL of phenol:chloroform:isoamyl alcohol (Sigma; pH 7-8) before the stool was added. High liquid content of stool resulted in increased sample volume after bead beating resulting in our scaling up of reagent volumes accordingly after which the manufacturer’s instructions were followed.

### Whole-genome sequencing

Whole-genome sequencing of *V. cholerae* and ICP1 isolates during the surveillance period was performed in our previous study (LeGault et al., 2021b). Briefly, DNA from *V. cholerae* and phages was extracted using commercially available kits (Qiagen DNeasy Blood and Tissue or Monarch genomic DNA purification kit (New England BioLabs). For preparation of phage DNA, phage stocks were treated with DNase at 37°C for 30 minutes before heat inactivation prior to DNA extraction following the manufacturer’s instructions. Libraries for Illumina sequencing were prepared with New England BioLabs’s Ultra II DNA or Ultra II FS DNA prep kits or performed by the Microbial Genome Sequencing Center. Sequencing (150 × 150 paired end) was performed by the QB3 Genomics Core at the University of California, Berkeley, or by the Microbial Genome Sequencing Center.

### CRISPR/Odn and PLE PCR screening

To detect ICP1-encoded anti-PLE loci from patient stool, total DNA from stool was used as a template for PCR using the primers listed in Table 1. Colony PCR using the primers listed in Table 1 was performed on individual *V. cholerae* colonies to screen for the presence of PLE1 amongst isolates recovered from the same patient sample.

### Genome database and PLE discovery

A database of publicly available genomes was generated by downloading all de-duplicated assemblies from the NCBI *V. cholerae* genome directory (https://www.ncbi.nlm.nih.gov/genome/browse/#!/prokaryotes/505/) and by a literature search for raw sequence reads available on the NCBI Sequence Read Archive (https://www.ncbi.nlm.nih.gov/sra). These raw reads were subsequently downloaded in fastq format and assembled with spades v3.14.0 using default settings. This is a similar, but expanded, dataset from previous work (LeGault et al., 2021b). Each of the five previously known PLEs were used as BLASTn queries against the completed *V. cholerae* database with default cutoffs. This was used to both identify PLE(+) genomes as well as discover similar sequences that could be potential new PLEs. These putative new PLEs were manually curated with CLC Main Workbench 7 to ensure similar length, synteny and sequence similarity to known PLEs.

### PLE alignment and phylogeny

PLE fasta files were aligned with progressiveMauve (v. 2015-02-25) using default parameters as well as the ‘assume colinear genomes’ setting. Region color combinations were randomly chosen from the Viridis color palette for each combination of PLEs and the phylogenetic tree was generated from the guide_tree file.

### PLE integration sites

The first 100 base pairs of the genomic regions immediately flanking every PLE positive *V. cholerae* isolate in the *V. cholerae* genome database were extracted and grouped by those that matched exactly. Only PLEs with a uniquely matched set of left and right regions flanking the integration site were considered (i.e. no left flank sequence occurred with more than one right flank sequence). This ensured that integration site differences were not due to genomic variability or shuffling within the superintegron of each PLE positive *V. cholerae* isolate. Those PLEs that satisfied these requirements were grouped by identical flanks pairs (i.e. integration sites) and plotted.

### Generation of bacterial and phage mutants

PCR constructs for bacterial mutations of interest were made through splicing by overlap extension and introduced by natural transformation ((Dalia et al., 2014)). To generate a PLE10(+) strain in the *V. cholerae* E7946 background PLE transduction was performed with magnesium as previously described (Netter et al., 2021). Phage mutants were constructed using CRISPR-Cas engineering (Box et al., 2016). All mutant strains were verified with Sanger sequencing over the region of interest.

### Assays for PLE’s response to ICP1 infection

Circularization of PLE upon infection was completed as previously described (McKitterick and Seed, 2018) with slight modifications. Briefly, *V. cholerae* strains were grown to OD600 ∼0.3 at which point an uninfected sample was taken, then the remaining culture was infected with ICP1_2006_E ΔCRISPR Δ*cas2-3* at a multiplicity of infection (MOI) of 2.5 for 20 minutes at 37°C. The uninfected and infected samples were then boiled for 10 minutes and 2 µL was used as a template for PCR using primers to detect the circularized PLE listed in Table 1. Replication of PLE upon ICP1 infection was measured as previously described (Barth et al., 2020; O’Hara et al., 2017). Briefly, 2mL cultures of *V. cholerae* were grown to an OD600 ∼0.3 before being infected with ICP1_2006_E ΔCRISPR Δ*cas2-3* (or the Δ*helA* derivative) at an MOI of 2.5. Immediately before the phage addition, 100 µL of culture was boiled for the T0 sample. Cultures were returned to the incubator. After 25 minutes (or 30 minutes where a plasmid construct was present), a 100 µL of infected culture was removed and boiled for the T25 sample. Serial dilutions of boiled samples were then used as the template for qPCR (primers listed in Table 1) and fold change in replication was determined as the amount of DNA in the T25 sample relative to the T0 sample. All samples were run in biological triplicates and technical duplicates. Lysis kinetics were determined a previously described (Hays and Seed, 2020). Briefly, *V. cholerae* strains were grown to OD600 = 0.3 in 2 mL cultures. Culture (150 uL) was then added to a 96-well plate containing phage. Kinetics were then recorded by the SpectraMax i3x (Molecular Devices) via measurements of OD600 every two minutes. Over the course of the run, the plate was incubated at 37°C and was shaken for 1 minute between reads. PLE transduction was performed with magnesium as previously described (Netter et al., 2021).

### Phage spot assays

Mid-log *V. cholerae* was added to 0.5% molten LB agar and poured on a solid agar plate and allowed to solidify. Ten-fold dilutions of phage were overlaid in 3 µL spots. After spots were dry, plates were incubated at 37°C for ∼6 hours prior to imaging. Images are representative of three independent experiments.

### *V. cholerae* phylogeny

A cladogram of 148 Vibrio cholerae isolates was constructed based on 532 single nucleotide variations (SNVs). These variable locations were determined by aligning the assembled sequence of each isolate to the reference strain N16961 using MUMmer (v. 3.0). Only nucleotide locations that were present in all 148 isolates as well as the reference were considered for analysis. The binary matrix of SNVs were clustered with R using R’s hclust package (v. 1.2.3). The SXT ICE present in each isolate was determined previously by whole-genome sequencing (LeGault et al., 2021b).

### Quantification and statistical analysis

For qPCR and transduction assays data each individual independent biological replicate is shown and the height of the bar indicates the average. For lysis curves, the shaded area indicates the standard deviation of the average fold change from three independent biological replicates. Spot plates and agarose gels are representative of at least three independent experiments.

