## Supplemental figures S1-S3 for "Evolutionary sweeps of subviral parasites and their phage host bring unique parasite variants and disappearance of a phage CRISPR-Cas system"

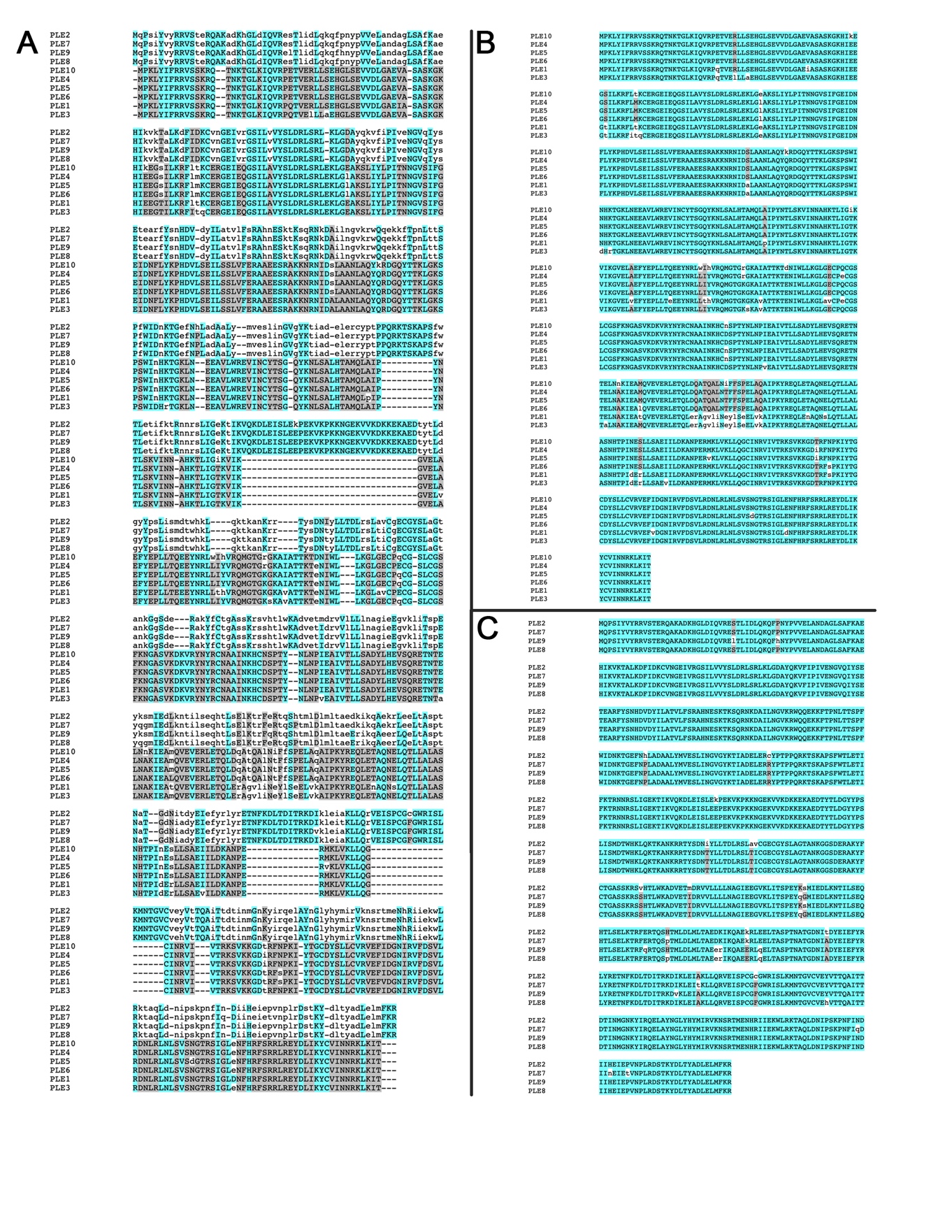

**Figure S1: Alignment of PLE integrases**

Coloring is determined by average BLOSUM62 score of pairs of letters in each column: light blue ≥ 1 or light gray ≥ 0.2. Otherwise no color. A) Alignment of integrases from all PLEs. B) Alignment of only PLE1-type integrases. C) Alignment of only PLE2-type integrases.

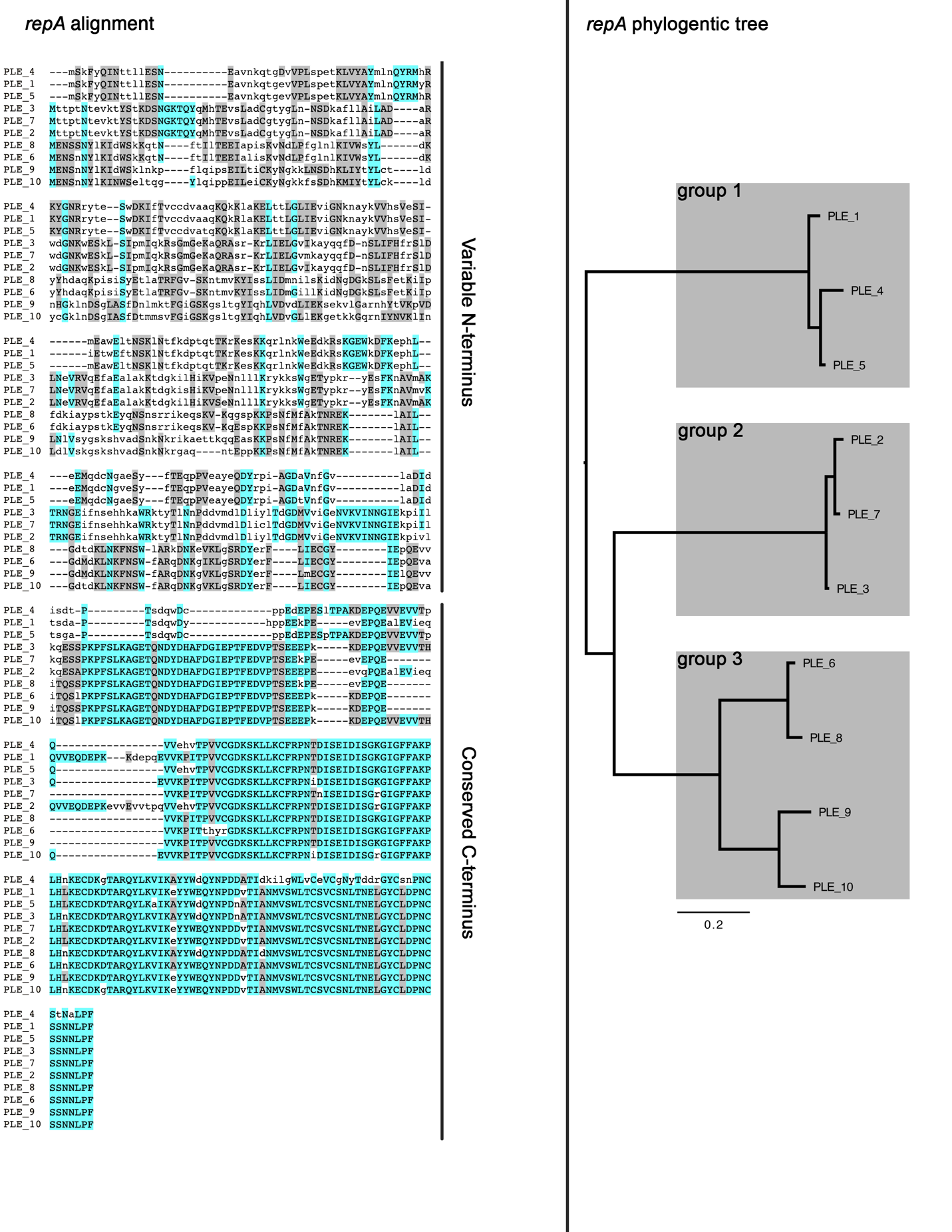

**Figure S2: RepA alignment and phylogenetic distance**

Alignment of translated *repA* genes with the same coloring as in Figure S1 (left). Phylogenetic tree of RepA amino acid alignment with major groups highlighted (right).

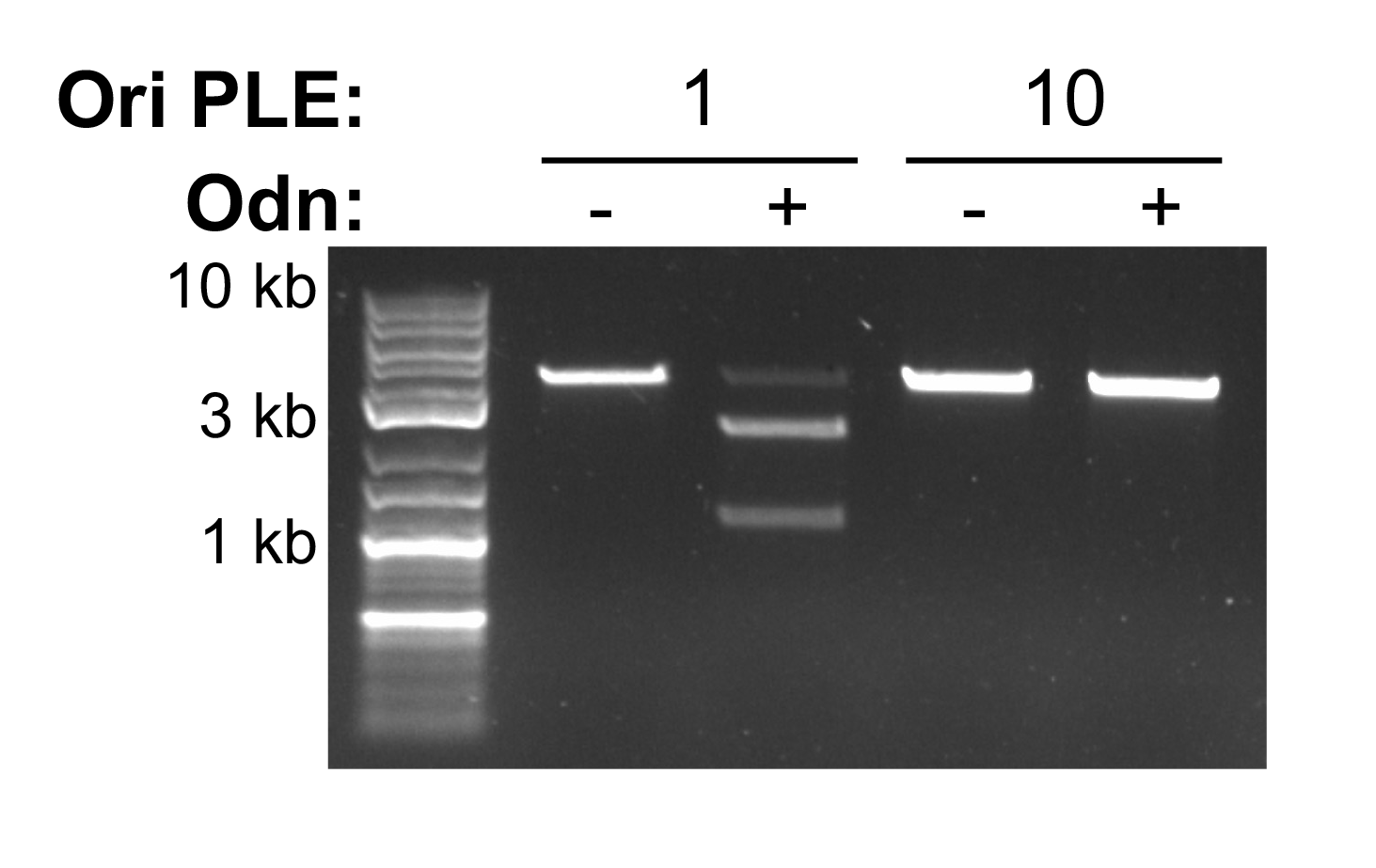

**Figure S3:** **The anti-PLE ICP1-encoded origin-directed nuclease (Odn) does not show activity against PLE10**.

Nuclease assay showing the integrity of a PCR product amplified from the noncoding region containing the ori from the PLE variant indicated (numbers) treated with (+) and without (–) 500 nM of purified Odn. Shown is representative of nuclease assays performed in triplicate.
